# Chromatin-lamin B1 interaction promotes genomic compartmentalization and constrains chromatin dynamics

**DOI:** 10.1101/601849

**Authors:** Lei Chang, Mengfan Li, Shipeng Shao, Boxin Xue, Yingping Hou, Yiwen Zhang, Ruifeng Li, Cheng Li, Yujie Sun

## Abstract

The eukaryotic genome is folded into higher-order conformation accompanied with constrained dynamics for coordinated genome functions. However, the molecular machinery underlying these hierarchically organized chromatin architecture and dynamics remains poorly understood. Here by combining imaging and Hi-C sequencing, we studied the role of lamin B1 in chromatin architecture and dynamics. We found that lamin B1 depletion leads to chromatin redistribution and decompaction. Consequently, the inter-chromosomal interactions and overlap between chromosome territories are increased. Moreover, Hi-C data revealed that lamin B1 is required for the integrity and segregation of chromatin compartments but not for the topologically associating domains (TADs). We further proved that depletion of lamin B1 leads to increased chromatin dynamics, owing to chromatin decompaction and redistribution toward nuclear interior. Taken together, our data suggest that chromatin-lamin B1 interactions promote chromosomal territory segregation and genomic compartmentalization, and confine chromatin dynamics, supporting its crucial role in chromatin higher-order structure and dynamics.

## Introduction

Chromatin in the interphase nucleus of eukaryotic cells are highly compartmentalized and structured. Owing to technological breakthroughs in imaging(1–3) and sequencing(4–9), chromatin higher-order structure has been increasingly studied over the last decade. Hierarchical chromatin architecture is composed of loops, TADs, active and inactive A/B compartments, and chromosome territories, in increasing scales. A number of architectural proteins and molecular machineries governing chromatin organization and dynamics have been identified. For instance, CTCF(10) and cohesin(11–14) are partly responsible for the formation and maintenance of chromatin loops and TADs. Nevertheless, CTCF does not impact higher-order genomic compartmentalization and cohesin even limits segregation of A/B compartments(15). Until now, the mechanisms that underlie the insulation and distribution of A/B compartments and chromosome territories remain elusive.

Microscopy and chromosome conformation capture techniques provide complementary insights into chromatin higher-order structure and sub-nuclear chromatin spatial distribution. Genomic regions that belong to A-compartments identified by Hi-C are gene rich, enriched with euchromatin histone markers and transcriptionally active(7, 16). Microscopy reveals that transcriptionally active euchromatic loci prefer to localize in nuclear interior(17). On the contrary, B-compartments identified by Hi-C are gene poor, enriched with heterochromatin markers, and frequently associated with the nuclear lamina(7, 16). These findings are concordant with imaging results that transcriptionally inactive heterochromatin is mainly found near nuclear periphery and nucleoli(17). In addition, chromosome territories also show nuclear location preference, in which gene-rich chromosomes are generally situated towards the interior and gene-poor chromosomes towards the periphery of the nucleus(18). Such spatial correlations make nuclear lamina a plausible candidate that contributes to the segregation and sub-nuclear distribution of A/B compartments and chromosome territories(19–21). Despite these insights, whether lamina proteins are responsible for the segregation and localization of higher-order chromatin structure remains unknown.

Besides the hierarchical structure, motion is another basic physical property of chromatin. Live cell fluorescence imaging in the nucleus can provide abundant information on the dynamics of whole chromosomes, chromosomal loci, transcription factors etc.(22). For instance, analyzing the motion of chromosomal loci embedded within viscoelastic nucleoplasm is a valuable technique for extracting micro-environmental properties and obtaining information of genomic organization and functions. Actually, the diffusion dynamics of genomic loci are not mere Brownian motion driven by thermal fluctuation but are often convoluted with active processes such as transcription and DNA repair(23–27). So far, most studies on genomic loci dynamics have been carried out in bacteria(28, 29) and yeast(30–33). In yeast, chromatin dynamics appear to be determined by nuclear constraints. In particular, the telomeres and centromeres are known to be tethered to nuclear envelope, which is suggested to contribute to chromosome territory segregation(34). For mammalian cells, previous studies have shown that chromatin motion is a constrained diffusion process that is associated with nuclear localization(35), and loss of lamin A increases chromatin dynamics in the nuclear interior as well as nuclear periphery(36). However, it is unclear how this sub-diffusion is related to chromatin higher-order structure, and it is not known whether tethering between the nuclear envelop and chromatin functions similarly as in yeast.

Nuclear lamina consists of many protein complexes, and lamins are the main components of nuclear lamina in most mammalian cells and can be classified into A- and B-type lamins. Lamin A and C are the most common A-type lamins and are splice variants of the same gene, while B-type lamins, B1 and B2, are the products of two different genes(37). Lamin B1 mainly localizes at the nuclear periphery, while A-type lamins are also found in the nucleoplasm(38). DamID of lamin B1 has revealed many nuclear lamina-associated genomic regions named lamina-associated domains (LADs). Typically, a mammalian genome contains 1100–1400 LADs and 71 % of the genome has conserved relationship with lamina across different species, including 33% of constitutive LAD (cLAD) and 38 % of constitutive inter-LAD (ciLAD)(39). Interestingly, lamin B1 has structural domains that directly bind to DNA or histone and indirectly interact with chromatin through LEM [LAP2 (lamina-associated polypeptide 2)/emerin/MAN1] domain-containing proteins. Thus, lamin B1 can potentially provide anchors for chromatin to regulate its position, higher-order structure and dynamics. Furthermore, lamina-chromatin junction mediates the transduction of mechanical signals from cytoskeleton to chromatin, possibly providing means for chromatin structures to respond to mechanical forces(40, 41).

In this study, we hypothesized that chromatin-lamina interactions in mammalian cells function in chromatin higher-order structure and dynamics. We applied a combination of imaging and sequencing techniques to characterize the role of lamin B1 in chromatin architecture and dynamics in human breast tumor cells. We found that lamin B1 is required for segregation of chromosome territories and A/B compartments, but does not affect TAD formation. Furthermore, depletion of lamin B1 or disruption of interaction between DNA and lamin B1 can increase genomic loci dynamics, owing to chromosome decompaction and redistribution toward nucleoplasm. Taken together, our data suggested that interactions between lamin B1 and chromatin greatly contribute to chromatin compartmentalization, compactness, spatial distribution and dynamics.

## Results

### Lamin B1 depletion leads to chromatin redistribution and decompaction

To explore the potential role of lamin B1 in nuclear chromatin organization, we first investigated the subnuclear distribution of lamin B1 using super-resolution (STORM) imaging. Lamin B1 was found to be almost exclusively located at the nuclear periphery (fig. S1A), in contrast to A-type lamins which were located at both nuclear periphery and nucleoplasm(36, 38). It was previously reported that lamin B1 interacts with chromatin directly or via adaptor proteins(42). We then created a *LMNB1* (lamin B1 encoding gene)-knockout MDA-MB-231 breast cancer cell line using the CRISPR-Cas9 genome editing tool. Proper knockout of lamin B1 was confirmed by western blot and immunofluorescence (fig. S1, A and B). Importantly, no apparent change of cell cycle was detected in lamin B1-KO cells (fig. S1D), eliminating the possibility that alterations of nuclear organization are due to biased cell cycle.

We reasoned that if the anchorage of chromatin to nuclear periphery is mediated by the interaction with lamin B1, the loss of lamin B1 can lead to changes in distribution and compaction of chromatin in the nucleus. To investigate the effect of lamin B1 on chromatin spatial localization and compaction at the single chromosome level, we performed chromosome painting for chromosomes 2 and 18 using FISH probes. Chromosomes 2 and 18 were chosen to represent chromosomes that are localized relatively near the nuclear periphery and the nuclear interior, respectively. Previous studies described that gene-poor chromosome 18 located toward the nuclear periphery and gene-dense chromosome 19 in the nuclear interior(43, 44). However, Cremer et al. reported that in seven of eight cancer cell lines, chromosome 18 located more internally than chromosome 19(45). Considering the breast cancer cell line we used, we chose chromosome 18 to represent nuclear interior localized chromosome rather than chromosome 19. In lamin B1-KO cells, chromosome 2 became significantly more centrally located while the position of chromosome 18 remained at the nuclear interior (Fig. 1, A and B). In addition, compared with wild type cells, the volume of both chromosomes were significantly increased in lamin B1-KO cells (Fig. 1, A and C). This expansion of chromosome territories upon lamin B1 depletion is not due to nuclear volume expansion (fig. S1E). These findings indicate that the nuclear location and volume of individual chromosomes are affected in lamin B1-KO cells, and suggest that loss of lamin B1 leads to redistribution and decompaction of the chromatin.

**Fig. 1.**
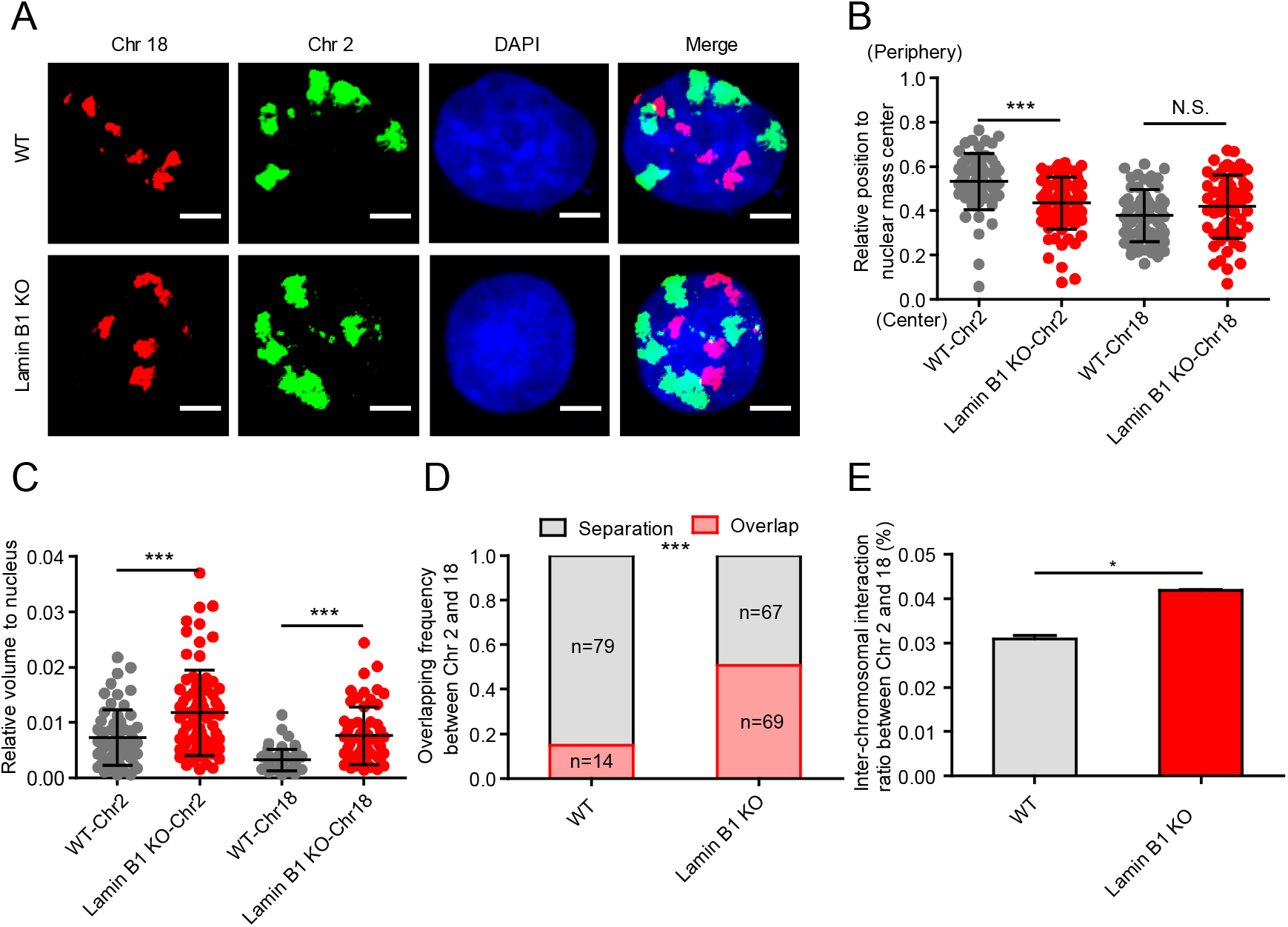
Lamin B1 regulates chromatin sub-nuclear localization and global compaction. (A) Two representative three dimensional (3D)-projection chromosome painting images of chromosome 2 and 18. Green: FISH signal of chromosome 2. Red: FISH signal of chromosome 18. Blue: DAPI staining. The maximum intensity projections of nuclear Z stacks are displayed. Scale bars, 5 µm.
(B) Quantification of the nuclear localization of chromosomes based on their relative distances from the chromosome mass center to the nuclear mass center. This distance is normalized by the cubic root of the nuclear volume. Mean ± standard deviation (SD). *** p < 0.001, Mann–Whitney test. 3 independent experiments.
(C) Quantification of the volumes occupied by chromosome 2 and 18 relative to the nuclear volume. Chromosomes in lamin B1-KO cells show significantly larger relative volumes. Mean ± SD. *** p < 0.001, Mann–Whitney test. 3 independent experiments.
(D) Quantification of the overlap frequency between chromosome 2 and chromosome 18 territories. The ratio of cells presenting territory interaction between chromosome 2 and chromosome 18 in wild type (WT) cells is significantly smaller than that in lamin B1-KO cells. *** p < 0.001, Fisher’s exact test. 3 independent experiments.
(E) Trans-interaction ratios of chromosome 2 and 18 in two WT replicates and lamin B1-KO replicates. Interaction numbers of chromosome 2 and 18 are normalized by the total interactions of the whole genome in each sample. Mean ± standard error (SE). * p<0.05, t-test.

### Lamin B1 depletion reduces the segregation of chromosome territories and A/B compartments

Changes in location and volume of chromosomes may affect the territories between chromosomes. Indeed, along with the redistribution and decompaction of chromatin, more than 50% of lamin B1-KO cells showed overlap between the territories of chromosomes 2 and 18, compared with 15.1% in wild type cells (Fig. 1, A and D). This large-scale reorganization of chromosome territories promoted us to investigate the role of lamin B1 in genome architecture using in situ Hi-C assay(46), which provides information about multiscale chromatin interaction maps including chromosome compartments and TADs (fig. S2, A and B, table S1). We first focused on inter-chromosomal interactions. In agreement with the FISH results (Fig. 1, A and D), Hi-C data showed higher inter-chromosomal interaction frequency between chromosomes 2 and 18 in lamin B1-KO cells (Fig. 1E and fig. S2C), although the interaction frequency between different chromosomes is much less than that within the same chromosome (table S1) as reported in previous studies(7, 47). The inter-chromosomal interaction ratio of all chromosomes also showed a significant increase in lamin B1-KO cells (Fig. 2, A and B). These results indicate that lamin B1 contributes to the segregation of chromosome territories.

**Fig. 2.**
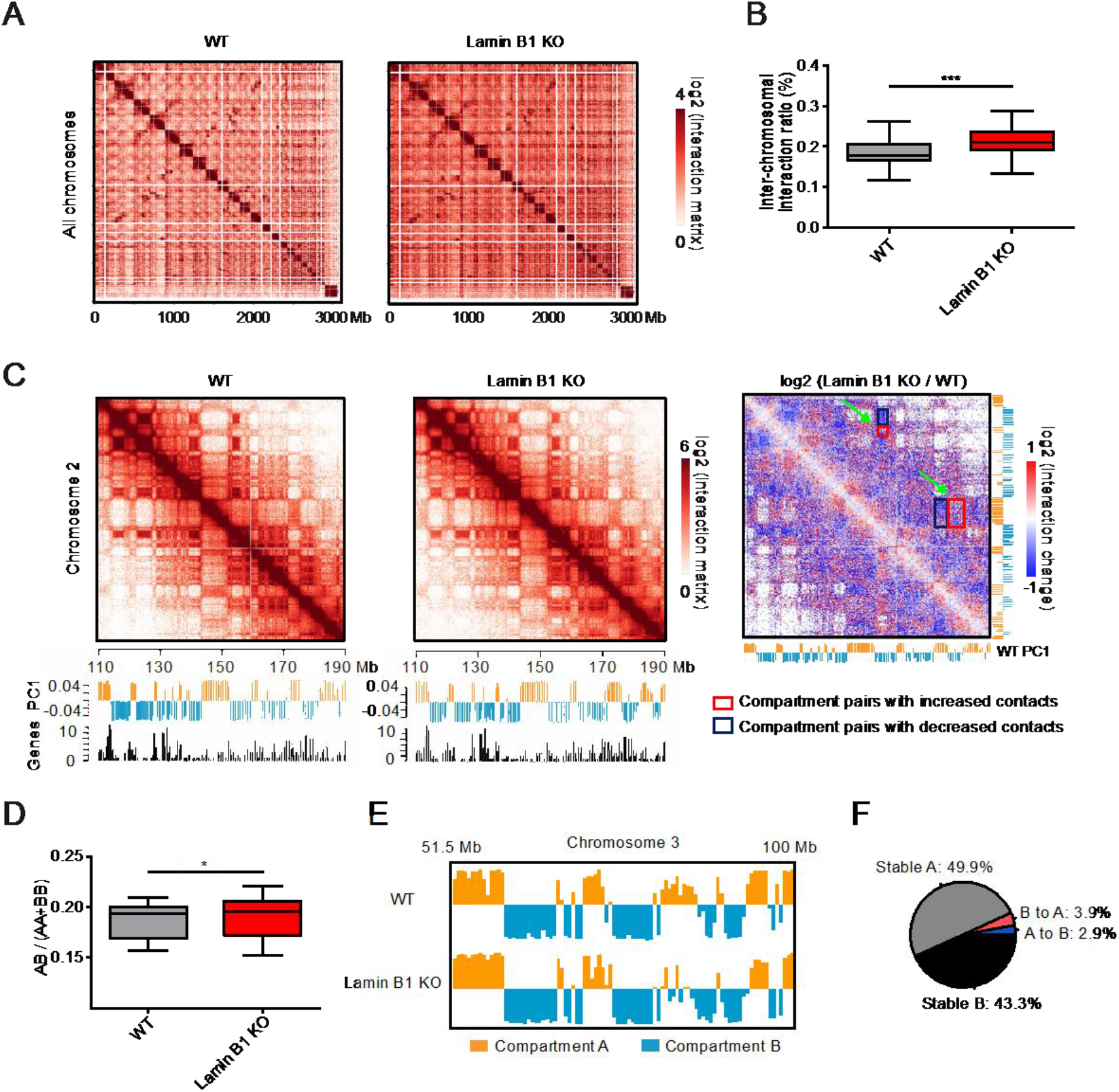
Lamin B1 depletion reduces the isolation of chromosome territories and A/B compartments. (A) Normalized Hi-C trans-interaction matrices for the whole chromosomes in WT and lamin B1-KO samples.
(B) Trans-interaction ratios of each chromosome in WT and lamin B1-KO cells. For each chromosome, trans-interaction ratio is the percentage of trans-interaction in total interaction of this chromosome. *** p<0.001, paired t-test. 2 biological repeats.
(C) Normalized Hi-C interaction matrices for chromosome 2 (110-190 Mb) in WT and lamin B1-KO cells, and differential matrices of genomic regions between WT and lamin B1-KO cells (resolution: 200 kb). Below the heatmaps are PC1 values and gene density plots. Orange represents compartment A, and blue represents compartment B. High gene density regions correlate with compartment A.
(D) Ratios of inter-compartment interactions (AB) and intra-compartment interactions (AA+BB) for each chromosome (X chromosome excluded) in WT and lamin B1-KO cells. *p<0.05, paired t-test. 2 biological repeats.
(E) Example of genomic region transition from A compartment in WT cells to B compartment in lamin B1-KO cells. Compartment A (orange, positive PC1 signal) and compartment B (blue, negative PC1 signal) distribution on chromosome 3 (51.5-100 Mb) in WT and lamin B1-KO cells.
(F) Genome-wide summary of genomic regions switching between A/B compartments in WT and lamin B1-KO cells. 2 biological repeats.

We next explored whether lamin B1-KO affects the organization of A and B compartments, which are defined using the first principal component (PC1) of Hi-C correlation matrices and correspond to different gene densities and transcriptional activities(7) (Fig. 2C). Using super-resolution imaging, Zhuang and colleagues have shown that adjacent A and B compartments are spatially separated from each other(2). Here, although the Hi-C contact maps of the wild type and lamin B1-KO cells displayed similar checkerboard patterns, the differential heatmap showed loss of intra-compartment interactions (interactions between the A-A or B-B compartment pairs) and gain of inter-compartment interactions (interactions between A-B compartment pairs) in lamin B1-KO cells (Fig. 2C). We asked whether this change influences the interactions between specific compartment types by computing the ratio of average interaction frequency between different classes of compartments (AB) versus that between the same classes of compartments (AA and BB) for each chromosome(48). These ratios showed significant increase in lamin B1-KO cells (Fig. 2D), suggesting a crucial role of lamin B1 in segregation of different compartment types. Moreover, 2.9% of genomic regions switched from A compartment in wild type cells to B compartment in lamin B1-KO cells, while 3.9% of genomic regions exhibited the opposite switching (Fig. 2, E and F). These percentages of compartment switching upon lamin B1 depletion were higher than those between replicates (fig. S2, D and E). These results indicate that lamin B1 contributes to the formation and segregation of different chromosomal compartment types.

### Lamin B1 is not required for TAD insulation

Within A/B compartments, chromatin is further packaged in the form of TADs, which are considered as the basic structural units of chromatin and are largely conserved between cell types and across species(49–51). We calculated insulation scores(51) for each 40 kb bin of the Hi-C normalized matrix, and the local minima of insulation scores indicated TAD boundaries. The contact maps and insulation scores of an example region on chromosome 10 showed similar TAD patterns in WT and lamin B1-KO cells (Fig. 3A). For the whole genome, insulation scores were highly correlated between WT replicates (Pearson correlation coefficient, r=0.984) or between lamin B1-KO replicates (r=0.987). Correlation between WT and lamin B1-KO samples (r=0.969) was only slightly lower than that between replicates (fig. S3A). Heatmaps showed that the distribution of insulation scores around TAD boundaries was similar between WT and lamin B1-KO cells (Fig. 3, B and C).

**Fig. 3.**
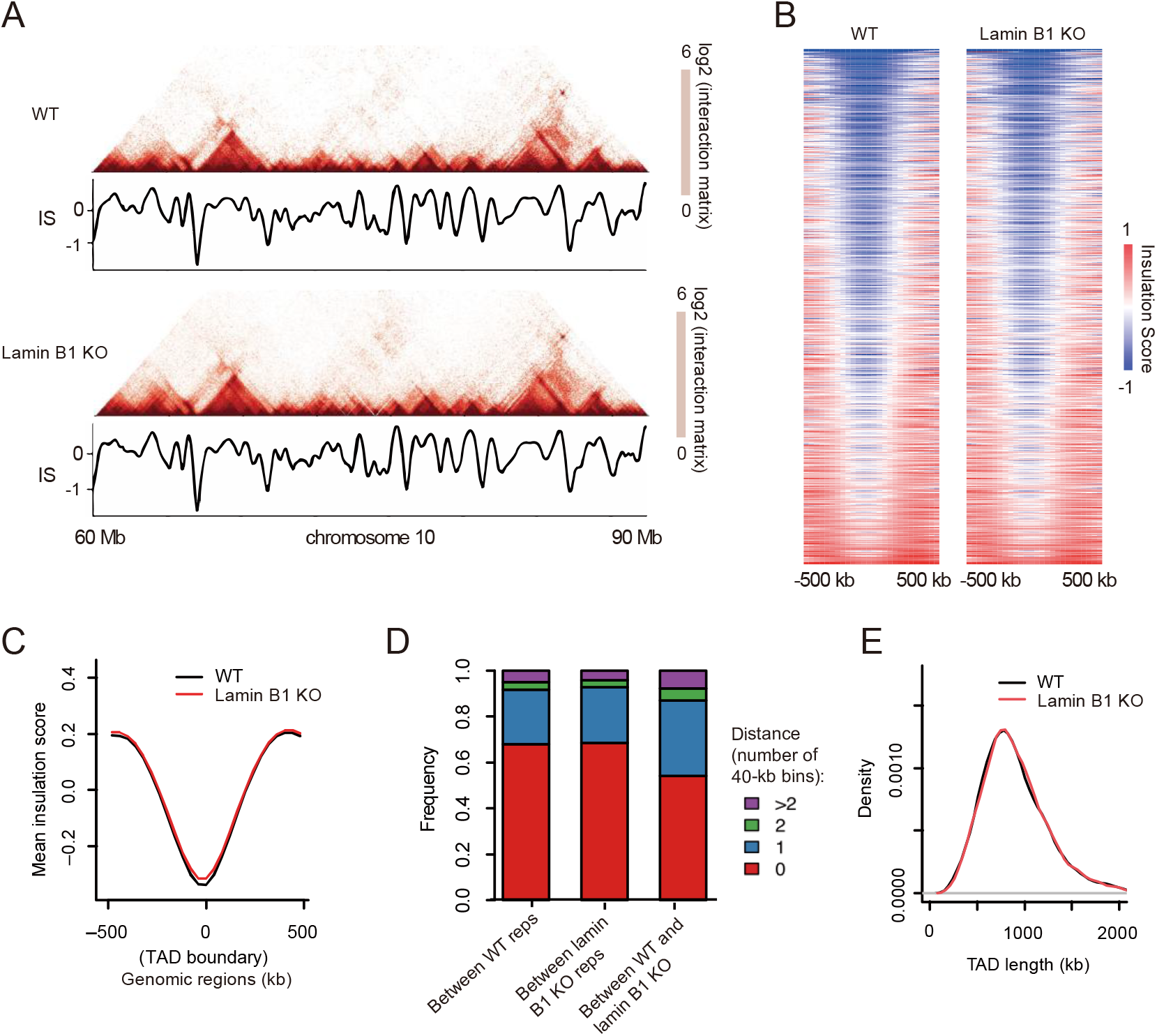
Lamin B1 is not required for TAD insulation. (A) Example of TAD pattern and insulation score distribution for chromosome 10 (60-90 Mb) in WT and lamin B1-KO cells.
(B) Heatmaps of insulation score around TAD boundaries in WT and lamin B1-KO cells. Heatmaps are organized according to the sum of insulation score around each boundary (±500 kb).
(C) Average insulation score distribution around TAD boundaries (±500 kb) in WT and lamin B1-KO cells. 2 biological repeats.
(D) Histogram of distance frequency of the most adjacent TAD boundary pairs between WT and lamin B1-KO cells, as well as between replicates of WT and lamin B1-KO-cells (40 kb bin).
(E) Distribution of TAD length in WT and lamin B1-KO cells. 2 biological repeats.

To investigate whether the TAD locations were changed upon lamin B1 depletion, each TAD boundary in WT cells was paired with the most adjacent TAD boundary in lamin B1-KO cells. We calculated the genomic distance between these paired TAD boundaries and observed that 87% of the TAD boundaries located within the same or adjoining 40 kb bins, and 92.3% of the TAD boundaries shifted by less than two 40 kb bins (Fig. 3D), comparable to these percentages (95% and 95.8%) between WT or lamin B1-KO replicates. The small number of TAD boundary pairs that are neither overlapping nor adjacent were due to random variation upon the calculation of insulation scores (fig. S3, B and C). As a result, WT and lamin B1-KO cells have almost overlapping TAD length distribution with median length of 840 kb (Fig. 3E). Furthermore, we calculated the TAD score, which is the ratio of intra-TAD interactions to overall cis-interactions, for each TAD, and found no difference between WT and lamin B1-KO cells (fig. S3, D and E), indicating similar TAD compactness for the two samples. Taken together, lamin B1 loss does not affect the organization of TAD structures.

### Lamin B1 depletion changes the location preference of genomic loci

To achieve high signal-to-noise ratio for precise localization and long-term imaging of genomic loci, we developed CRISPR-SunTag, a site-specific chromatin labeling and tracking system, in wild type and lamin B1-KO MDA-MB-231 cells (Fig. 4A and fig. S4, A and B). To quantitatively categorize the position of genomic loci, we divided the nuclear space into two regions of different transcription activities, i.e. nuclear periphery and nucleoplasm (Fig. 4B and see Materials and Methods for details). Genomic loci imaging showed that the same genomic loci could localize in different subnuclear regions (Fig. 4C) but did demonstrate location preferences (Fig. 4D). For instance, the 1 Mb genomic locus on chromosome 2 showed high percentage of nuclear peripheral localization (Fig. 4, C and D), consistent with the overall peripheral location preference of chromosome 2 (Fig. 1, A and B), while the 236 Mb locus on chromosome 2 showed low percentage of nuclear peripheral localization (Fig. 4, C and D). In contrast, genomic loci on chromosome 18 tended to distribute in the nucleoplasm (Fig. 4, C and D). In order to avoid measurement artifacts caused by projection from 3D to 2D, we compared the measured distances between loci and nuclear envelope or nucleoli in 2D images and 3D image stacks and obtained similar results (fig. S4, C and D). Thus, the overall location of genomic loci in the nucleus coincides with their corresponding chromosome localization, but different loci on the same chromosome have variable preferential subnuclear localization.

**Fig. 4.**
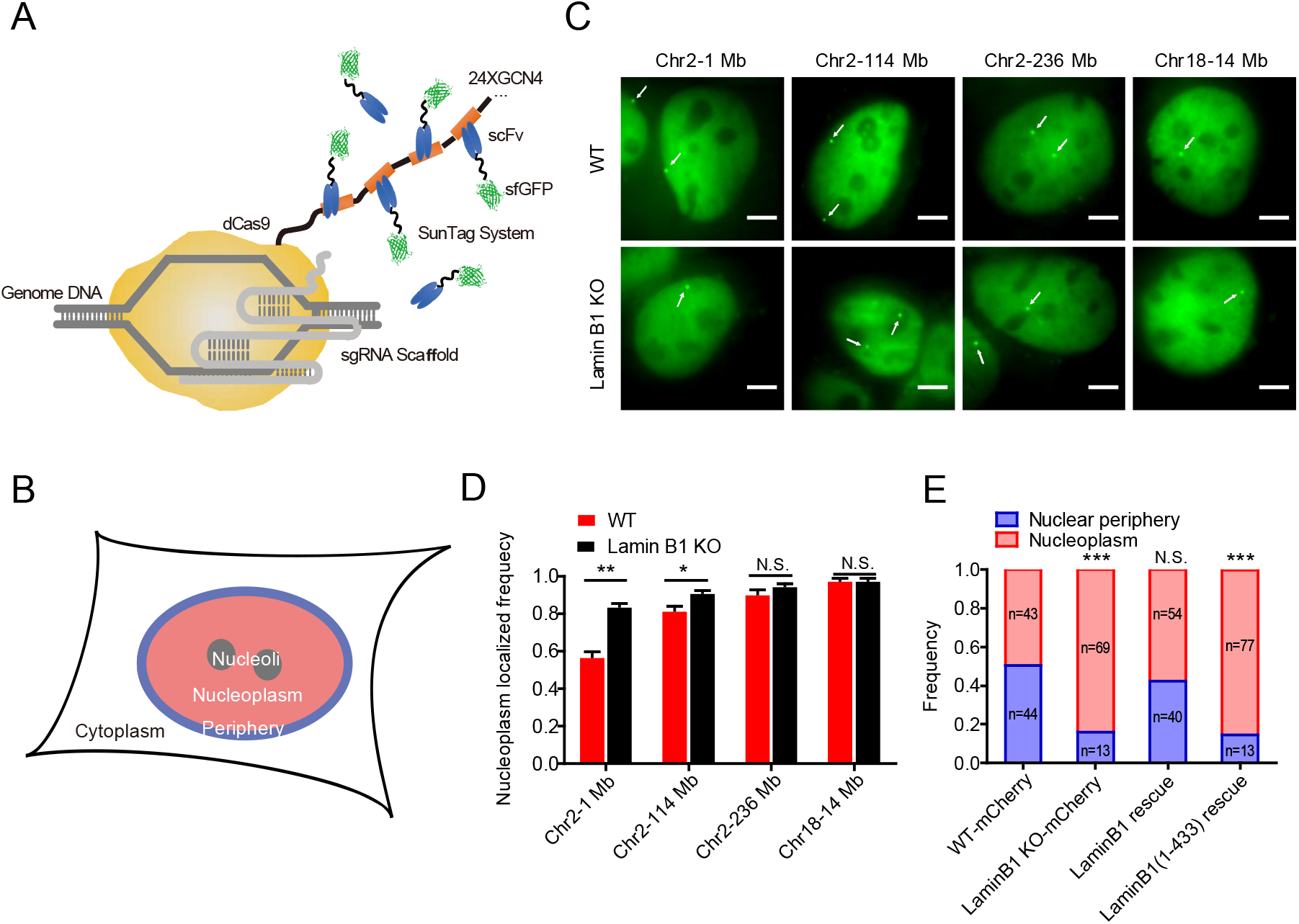
Lamin B1 depletion changes the location preference of genomic loci. (A) Schematic representation of CRISPR-SunTag, a labeling and signal amplification system including dCas9 fused with 24 tandem repeats of GCN4 peptide and a sfGFP-tagged single chain antibody (scFv) for GCN4 peptides. Using dCas9-(GCN4)_24x_ coexpressing with scFv-GCN4-sfGFP at minimal level, a single sgRNA can recruit as many as 24 fluorescent proteins to the target site.
(B) Each nucleus is divided into two compartments, nuclear periphery (blue) and nucleoplasm (pink, including nucleoli).
(C) CRISPR-SunTag labeling of chr2-1 Mb, chr2-114 Mb, chr2-238 Mb and chr18-14 Mb in WT and lamin B1-KO cells. The white arrows show signals of each loci. Scale bars, 5 µm.
(D) The nucleoplasm-localizing frequency of chr2-1 Mb (n=84), chr2-114 Mb (n=117), chr2-238 Mb (n=97) and chr18-14 Mb (n=80) in WT cells, as well as those of chr2-1 Mb (n=88), chr2-114 Mb (n=97), chr2-238 Mb (n=108) and chr18-14 Mb (n=85) in lamin B1-KO cells. ** p < 0.01, * p < 0.05, unpaired t test. 3 independent experiments.
(E) The subnuclear localization changes of 1Mb loci in chromosome 2 in mCherry expressing-WT cells (n=87), mCherry expressing-lamin B1-depleted cells (n=82), lamin B1-rescue cells (n=94) and lamin B1(1-433)-rescue cells (n=90). *** p < 0.001, Fisher’s exact test. 3 independent experiments.

Lamin B1 depletion showed minimal effect on the preferential subnuclear distribution of genomic loci on chromosome 18 but dramatically altered that on chromosome 2, especially the loci near the nuclear periphery (Fig. 4, C and D). For example, the percentage of the 1 Mb locus of chromosome 2 localized near the nuclear periphery was greatly decreased in lamin B1 depleted cells (Fig. 4, C to E and fig. S4, E and F). Lamin B1 may regulate the genomic loci distribution via direct/indirect binding interaction or spatial confinement of accumulative nuclear lamina proteins in nuclear periphery. To distinguish between these two possibilities, we constructed a plasmid expressing a lamin B1 truncation protein missing the Ig-like domain. The Ig-like domain is a conservative structure in lamin A/C and lamin B1(52), known as the motif that mediates the direct/indirect interaction between lamins and DNA(42, 53) (fig. S4G). In contrast to the exogenous full-length lamin B1 which could rescue the distribution preference of the 1 Mb locus on chromosome 2 in lamin B1-KO cells, the Ig-like domain-truncated lamin B1 failed to do so (Fig. 4E) even though it could still form the nuclear lamina (fig. S4H). These results suggest that the tethering between lamin B1 and chromatin is important for the subnuclear position of chromosomes and chromosomal loci.

### Loss of chromatin-lamin B1 interaction increases chromatin mobility

Chromatin structures and transcriptional activities are intrinsically associated with its dynamic motion(24, 27, 54). To further explore the influences of chromatin-lamin B1 interaction on the chromatin, we measured the chromatin dynamics in wild type and lamin B1-KO cells. Three genomic loci consisting of telomeres, a locus localized at 1 Mb of chromosome 2 and a locus at 14 Mb of chromosome 18, were labeled and successively tracked in a short range of time scales (from 0.05 to 120 s) to minimize the artifacts caused by cell deformation, migration or nucleus rotation (Movie). A customized tracking package U-track(55) was used to extract the trajectories and mean square displacement (MSD) of the loci. The data revealed that depletion of lamin B1 significantly increased the chromatin dynamics compared with the slow anomalous diffusion in wild type cells of all three loci (Fig. 5, A and B). Moreover, expressing exogenous lamin B1 in the knockout cell line restored the loci dynamics to the level comparable to wild type cells (Fig. 5, C and D), indicating that lamin B1 restricts chromatin dynamics. However, Ig-like domain-truncated lamin B1 was not able to restore the loci dynamics in lamin B1-KO cells to the wild type level as the full length lamin B1 did (Fig. 5, C and D), coinciding with the changes of loci localization (Fig. 4E).

**Fig. 5.**
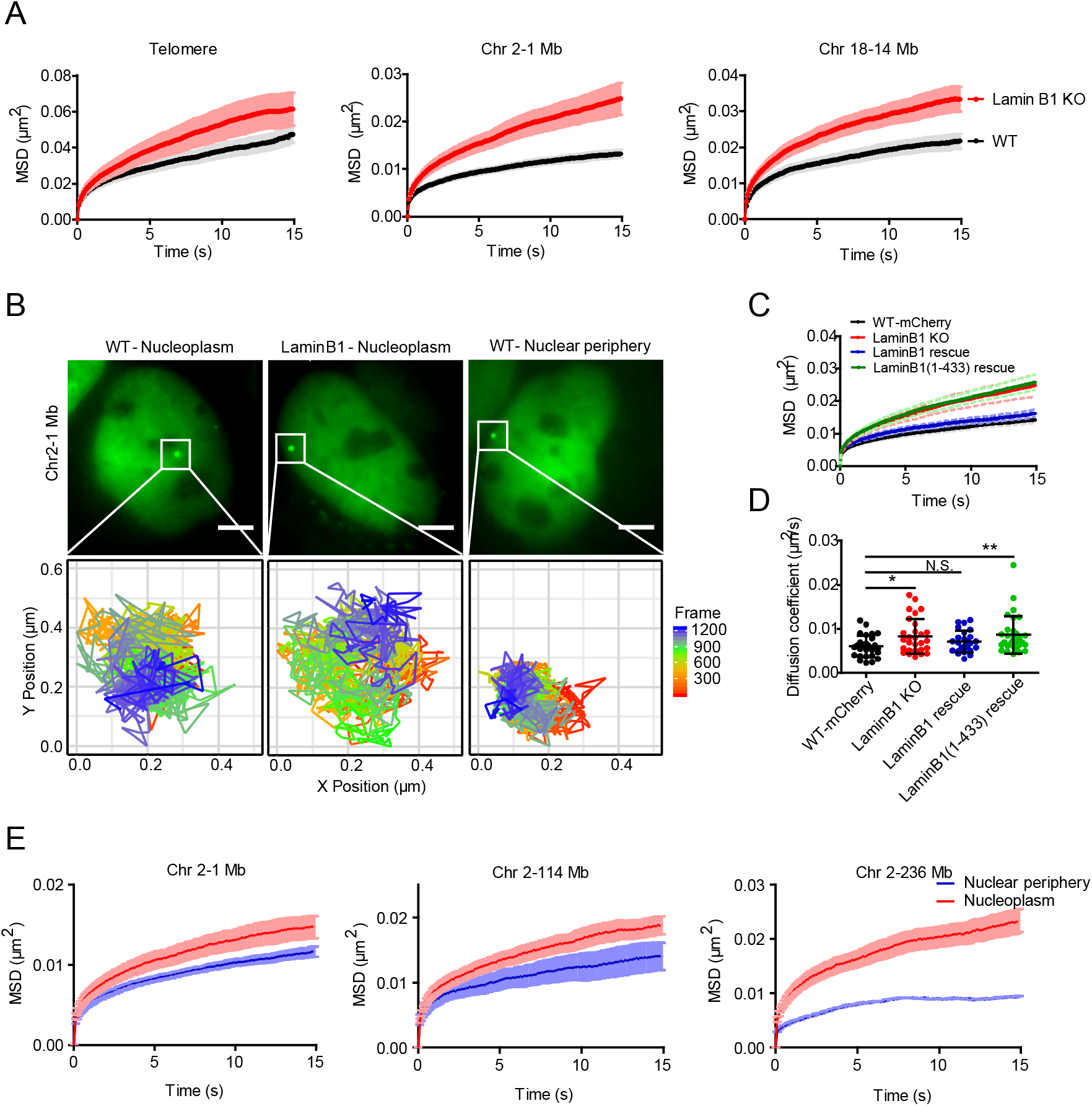
Loss of chromatin-lamin B1 interaction increases chromatin mobility. (A) MSD curve of telomeres in WT (n=100) and lamin B1-KO (n=86) cells. MSD curves of 1Mb loci on chromosome 2 in WT (n=27) and lamin B1-KO (n=29) cells. MSD curves of 14 Mb loci on chromosome 18 in WT (n=28) and lamin B1-KO (n=33) cells. Mean ± SE. 3 independent experiments.
(B) The tracking trajectories of labeled 1Mb loci on chromosome 2 in nucleoplasm of WT cells, nucleoplasm of lamin B1-KO cells and nuclear periphery of WT cells. Different colors of trajectories represent time lapse. Scale bars, 5 µm.
(C) MSD curve of 1Mb loci in WT (expressing mCherry, n=29), lamin B1-KO (n=29), lamin B1-rescue (n=25) and lamin B1(1-433)-rescue (n=30) cells. Mean ± SE. 3 independent experiments.
(D) The diffusion coefficient of 1Mb loci in WT (expressing mCherry, n=29), lamin B1-KO (n=29), lamin B1-rescue (n=25) and lamin B1(1-433)-rescue (n=30) cells. Mean ± SD. p < 0.05, ** p < 0.01, unpaired t test. 3 independent experiments.
(E) 3 genomic loci on chromosome 2 are tracked and assigned to nuclear periphery or nucleoplasm compartment, including 1 Mb loci (n=27), 114 Mb loci (n=30) and 236 Mb loci (n=18). Averaged MSD curves of these loci in the two compartments are calculated and displayed as mean ± SE. 3 independent experiments.

**Fig. 6.**
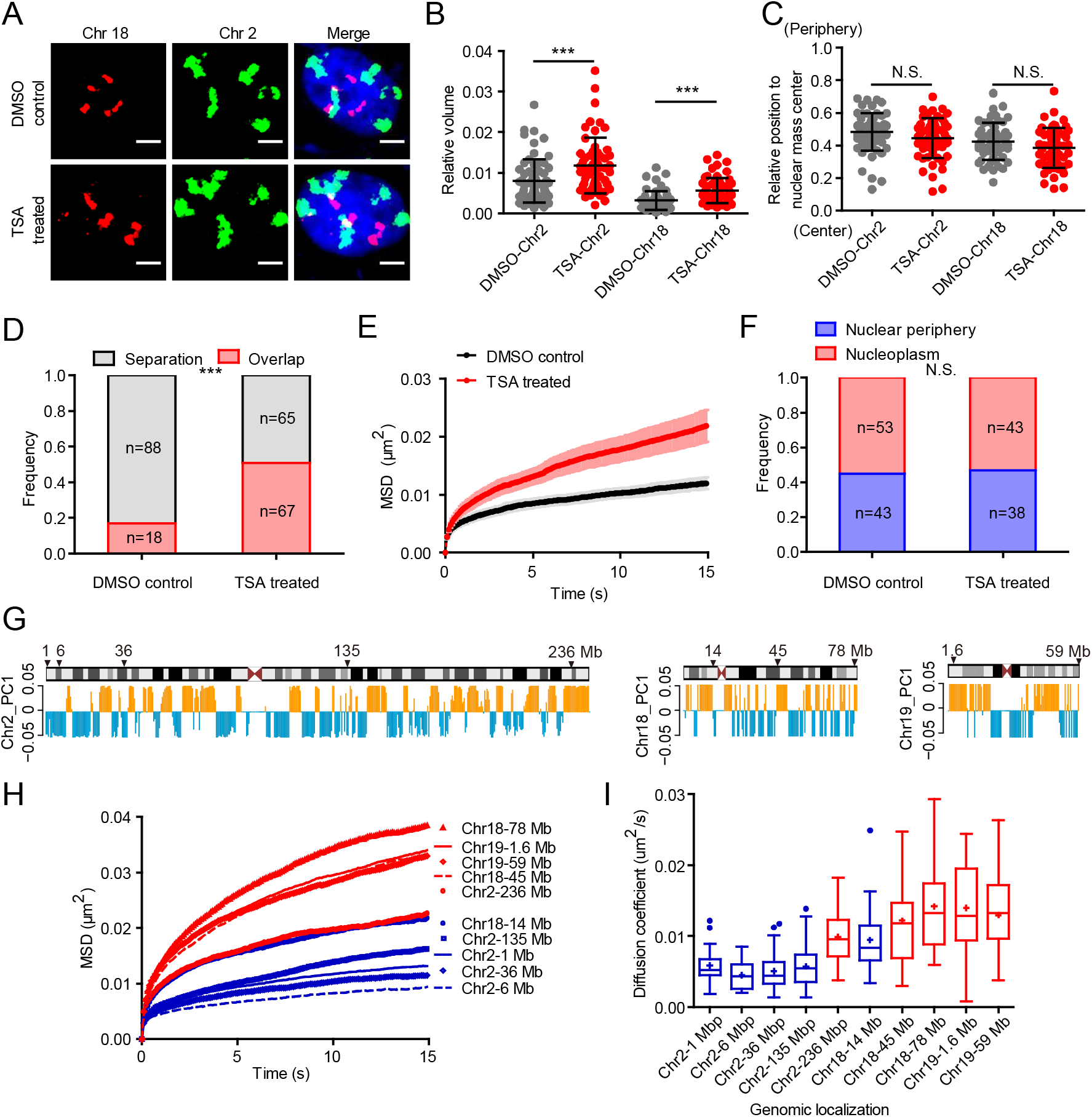
Global decompaction of chromatin contributes to chromatin dynamics increase and chromosome territories intermingling. (A) Representative 3D-projection chromosome painting images of chromosome 2 and 18 in DMSO-treated control cells and TSA-treated cells. Green: FISH signal of chromosome 2. Red: FISH signal of chromosome 18. Blue: DAPI staining. The maximum intensity projections of nuclear Z stacks are displayed. Scale bars, 5 µm.
(B) Quantification of the volumes occupied by chromosome 2 and 18 relative to the nuclear volume. Mean ± SD. *** p < 0.001, Mann–Whitney test. 3 independent experiments.
(C) Quantification of the nuclear localization of chromosomes based on their relative distances from the chromosome mass center to the nuclear mass center. This distance is normalized by the cubic root of the nuclear volume. Mann–Whitney test. 3 independent experiments.
(D) Quantification of the overlap frequency between chromosome 2 and chromosome 18 territories. The ratio of cells presenting territory interaction between chromosome 2 and chromosome 18 in control cells (n=106) is significantly smaller than TSA-treated cells (n=132). *** p < 0.001, Fisher’s exact test. 3 independent experiments.
(E) MSD curves of 1Mb loci on chromosome 2 in DMSO-treated control cells (n=25) and TSA-treated cells (n=26). Mean ± SE. 3 independent experiments.
(F) The spatial localization of 1 Mb loci on chromosome 2 in DMSO-treated control cells (n=96) and TSA-treated cells (n=81). Fisher’s exact test. 3 independent experiments.
(G) Schematic representation and PC1 values plots of chromosome 2, chromosome 18 and chromosome 19. Orange represents compartment A, and blue represents compartment B. Arrows indicate the genomic distribution of chosen loci.
(H) The averaged MSD curves and (**I**) The diffusion coefficient distribution of 10 genomic loci on chromosome 2, chromosome 18 and chromosome 19. Red indicates loci belonging to A compartment and blue indicates loci belonging to B compartment. 3 independent experiments.

We next investigated how chromatin-lamin B1 interaction constrains chromatin dynamics. We speculated that the redistribution of genomic loci in lamin B1-depleted cells contributes to increased chromatin mobility. The above results promoted us to examine whether the same locus in different subnuclear regions demonstrate different dynamic mobility. Indeed, all three loci on chromosome 2 were much less mobile when located in the nuclear periphery than in the nucleoplasm (Fig. 5, B and E), suggesting that the dynamics of each locus is primarily influenced by their nuclear spatial environment.

Furthermore, we found that the motion of the 1 Mb locus on chromosome 2 in both nuclear periphery and nucleoplasm became more active upon the depletion of lamin B1 (fig. S5A). Besides, the 14 Mb locus on chromosome 18, which only had nucleoplasm distribution, also showed increased mobility in nucleoplasm in lamin B1-KO cells (Fig. 5A). These results indicate that lamin B1 restrains the mobility of genomic loci in both nuclear periphery and nucleoplasm, not in line with the nuclear lamina distribution of lamin B1. Thus, our data indicate that lamin B1 also constrains chromatin dynamics through other ways, especially in the nucleoplasm.

### Chromatin decompaction mediates the effect of lamin B1 depletion on chromatin dynamics

Given the finding that lamin B1 depletion leads to chromatin decompaction (Fig. 1, A and C), we next examined whether the increased chromatin dynamics upon loss of lamin B1 was due to chromatin decompaction. We treated wild type cells with Trichostatin A (TSA) which can inhibit histone deacetylase enzyme and lead to genome-wide decondensation of chromatin in both nuclear interior and periphery(56). To confirm the effect of chromatin decompaction on chromosome spatial organization, we applied chromosome painting in TSA-treated cells. We found that the relative volume of chromosomes increased significantly compared with that in control cells (Fig. 6, A and B), but different from lamin B1 depletion which also altered the position of chromosomes (Fig. 1, A and B), TSA treatment did not change the radial distribution of chromosome territories (Fig. 6, A and C). The overlap between chromosome 2 and 18 was also consequently increased in TSA-treated cells compared with control cells (Fig. 6, A and D). We then measured the dynamic mobility of chromosomal loci and found that TSA treatment indeed promoted the dynamic mobility of chromosomal loci both near the nuclear periphery and within the nucleoplasm (Fig. 6E and fig. S5B). Importantly, different from lamin B1 depletion, the sub-nuclear distribution of the loci did not change in TSA-treated cells compared with DMSO-treated control cells (Fig. 6F). These results suggest that chromatin compaction is key for chromatin dynamics.

To further explore the relationship between chromatin dynamics and chromatin compaction state, we chose 10 genomic loci on chromosome 2, 18 and 19 with 5 in A compartments and 5 in B compartments (Fig. 6G). Tracking of the loci showed that the 5 loci belonging to A compartments (red) were more mobile than the 5 loci in B compartments (blue) (Fig. 6, H and I), in line with the fact that A compartments are generally less compact than B compartments. We also found that four genomic loci belonging to A compartments in chromosome 18 and 19 had similar dynamic properties and all of them localized in nucleoplasm (Fig. 6, H and I).

## Discussion

Along with chromatin decompaction and relocation, the motion of genomic loci became more active in lamin B1-depleted cells. In control cells, mobility of the same chromosomal locus was found to be correlated with its subnuclear location in that genomic sites close to lamina were generally less mobile than those localized in nuclear interior. These two lines of evidence suggest that chromatin dynamics are dependent on both chromatin compaction and loci location. However, when treating cells with TSA, which decompacts chromatin but does not change its sub-nuclear distribution, we also found global increase of chromatin mobility. Furthermore, in control cells, we consistently observed that chromosomal loci in less compact compartment A were of higher mobility than that in more compact compartment B, suggesting that chromatin compaction is more fundamental than subnuclear location in regulating chromatin dynamics. This observation is consistent with a recent theoretical work in which a model named MiChroM was proposed for the formation of chromosomal spatial compartments(54). MiChroM defines dynamically associated domains (DADs) in which the motions of genomic loci are correlated. DADs are often found to be aligned with the A/B chromatin-type annotation and another study proposed that the globally increased mobility of genomic loci may drive re-segregation at the chromatin compartment level via modifying MiChroM(57). This theoretical work is highly complementary with our experimental data, supporting the important role of chromatin dynamics in higher-order chromatin organization.

The roles of lamins played on chromatin organization are less understood. Recently, Leonid Mirny et al developed a polymer model of chromosomes to reconstruct chromatin sub-nuclear localization in inverted and conventional nuclei(58). They found that heterochromatin interactions with the lamina are essential for building conventional nuclear architecture. In our study, we found that interaction between lamin B1 and chromatin could drive the regulation of global chromatin structure (fig. S6). However, regarding the nature of lamin B1-chromatin interaction, it is unclear whether the direct binding or the confinement by the lamina meshwork mainly contributes to the regulation of chromatin structure and dynamics. Here our work has provided several lines of evidence to support the direct binding model. First, over-expressing Ig-like domain-truncated lamin B1, which can still form meshwork, was not able to rescue the phenotype of chromatin structure and dynamics caused by loss of lamin B1. Second, overexpressing lamin B1 did not alter the chromatin dynamics. Therefore, it is intriguing to speculate that the tethering and release of chromatin from lamina during each cell cycle might be an important process to organize genome architecture.

Previous studies have shown that other protein components in lamina besides lamin B1 also mediate its interactions with chromatin, including lamin A/C and LBR(59). Y. Garini et al reported that depletion of lamin A can increase chromatin dynamics(36). They emphasized that chromosomal inter-chain interactions formed by lamin A throughout the nucleus is critical for the maintenance of genome organization but did not focus on the tethering of chromatin with lamin A in the nuclear envelop. Interestingly, triple knockout of lamins did not show effects on overall TAD structure but altered TAD-TAD interactions in mESC(19), which is consistent with our results. However, lamins-KO mESCs had no significant differences in the volumes and surface areas of chromosome 1 and 13, which is different from previous studies and our results, indicating that lamins may play a stronger role in maintaining chromatin structure in differentiated cells than in ESCs(19, 60). Thus, the meshwork caging model they proposed may partially but not totally apply to differentiated cell types. Further studies are needed to identify more functional proteins in the lamina and their functional roles in chromatin structure and dynamics.

The nuclear matrix, which is hypothesized to provide a scaffold for chromatin attachment and organize global chromatin structure in the nucleus, is composed of inner and peripheral nuclear matrix. Recently, Hui Fan et al reported that the inner nuclear matrix protein HNRNPU/SAF-A is involved in 3D genome organization(61). We compared our lamin B1 data with their HNRNPU/SAF-A data and interestingly found that they contribute to chromatin organization in an opposite manner, implicating some fundamental coordinations between inner and peripheral nuclear matrix in regulation of chromatin structures. For instance, in contrast to our findings that depletion of lamin B1 promotes chromatin decompaction and relocalization from nuclear periphery to nucleoplasm, loss of HNRNPU promotes global condensation of chromatin and increases lamina-associated genomic regions. Moreover, genes enriched in cell adhesion are up-regulated in HNRNPU depleted cells but down-regulated in lamin B1 knockout cells (fig. S7). At the A/B compartment level, depletion of HNRNPU and lamin B1 both result in ∼10% transition between A/B compartments. Therefore, these two studies demonstrate that the inner and peripheral nuclear matrix, through anchoring of chromatin in the nucleoplasm and the nuclear envelop respectively, may offer a complementary, tug-of-war regulation of higher-order chromatin organization.

In this work, we combined imaging and Hi-C sequencing to study the role of lamin B1 in chromatin architecture and dynamics. We found that lamin B1 contributes to segregation of chromosome territories and A/B compartments, and loss of lamin B1 leads to chromatin decompaction and relocalization from nuclear periphery to nuclear interior. Besides, loss of lamin B1 changes the location preference of genomic loci and increases chromatin dynamics. Disruption of interactions between lamin B1 and DNA using truncated lamin B1 leads to similar effects on chromatin dynamics caused by loss of lamin B1. Dynamics of the same genomic locus is found to be correlated with its nuclear spatial environment in that genomic sites close to the lamina are generally less mobile than those localized in the nuclear interior. Our study demonstrates that it is chromatin decompaction that mediates the effect of lamin B1 depletion on chromatin dynamics. Furthermore, our study reveals that genomic loci in less compact compartment A are of higher mobility than those in more compact compartment B. Taken together, our work supports lamin B1 plays a crucial role in chromatin higher-order structure and chromatin dynamics.

## Materials and Methods

### Construction of sgRNA expression plasmids and SunTag PiggyBac plasmids

The mining process for repeats was similar as described recently(62). Briefly, the human genome sequence was downloaded from the UCSC genome browser (http://genome.ucsc.edu) with undetermined regions “Ns” replaced by randomly generated nucleotides “A”, “T”, “G”, or “C”. Then the sequence was input to the Tandem Repeat Finder bioinformatics tool (http://tandem.bu.edu/trf/trf.html) to identify the tandem repeats. Highly conserved repeats with little mutation, proper repeat unit length and repeat number were selected as candidates for live cell fluorescent labeling and imaging. sgRNA oligoes targeting each repeat were designed upstream of proto-spacer adjacent motif (PAM) sequence “NGG”. The oligoes of each sgRNA that target the repeat regions on human chromosomes were synthesized by Beijing Ruibo biotech (Beijing, China) with a 4 bp overhang flanking the sense and antisense strands. The sgRNAs targeting lamin B1 gene were designed by the online tool Optimized CRISPR Design (http://crispr.mit.edu) and candidates with the highest score were selected. The sgRNA expression vector for imaging was based on the psgRNA2.0 transient expression plasmid with an A-U flip and stem-loop extension (a gift from Prof. Wensheng Wei, Peking University), containing the ccdB screening gene between two BsmBI sites for inserting guide sequences into the sgRNAs. The sgRNA expression vector for editing was based on plasmid pX330-U6-Chimeric_BB-CBh-hSpCas9 (Addgene Plasmid # 42230), containing two BpiI restriction sites for inserting guide sequences into the sgRNAs. The targeting sgRNA expression plasmids were made by replacing the lethal gene ccdB with annealed oligo using Golden Gate cloning with enzyme BsmBI and T4 ligase (NEB). For the sequence of each sgRNA construct.

In order to construct a stable cell line, the NLS_SV40_-dCas9-3X NLS_SV40_-24X GCN4_-V4_-NLS_SV40_-P2A-BFP fragment was amplified by PCR from plasmid pHRdSV40-NLS-dCas9-24xGCN4_v4-NLS-P2A-BFP-dWPRE (Addgene Plasmid #60910) and then ligated into PiggyBac plasmid pB-TRE3G-BsmBI-EF1α-PuroR-P2A-rtTA by Golden Gate Assembly with enzyme BsmBI and T4 ligase (NEB). The ScFV-sfGFP-GB1-NLS_SV40_ fragment was amplified by PCR from plasmid pHR-scFv-GCN4-sfGFP-GB1-NLS-dWPRE (Addgene Plasmid # 60906) and then ligated into PiggyBac plasmid pB-TRE3G-BsmBI-EF1α-HygroR-P2A-rtTA by Golden Gate Assembly with enzyme BsmBI and T4 ligase (NEB).

### Cell culture, transfection and TSA treatment

Human cell line MDA-MB-231 cells were maintained in Dulbecco’s modified Eagle medium with high glucose (Lifetech). The medium contained 10% Fetal bovine serum (FBS) (Lifetech), and 1% of penicillin and streptomycin antibiotics (Lifetech). Cells were maintained at 37 °C and 5% CO_2_ in a humidified incubator. All plasmids were transfected with Chemifect (Beijing Fengrui Biotech, Beijing, China) in accordance with the manufacturer’s protocol. TSA (TricostatinA, Sigma-Aldrich) was eluted to 3 mM in DMSO. Cells were treated with 300 nM of TSA solution in complete growth medium for 24 hr before imaging experiments and the negative control sample was treated with DMSO.

### CRISPR-mediated lamin B1 gene knockout

In order to knockout lamin B1 genes, the cells were co-transfected with corresponding sgRNA and Cas9 chimeric plasmid and an empty mCherry expressing plasmid. At 48 h post transfection, cells were subjected to FACS to isolate mCherry positive single cell clone in 96-well plates. After incubation for about a month, genome of each grown clone was extracted and PCR-amplified with lamin B1-specific primer and sent for Sanger sequencing. Clones with indel were verified by Western Blot and immunofluorescence.

### Construction of the SunTag stable cell line

To construct the stable cell line, MDA-MB-231 cells were spread onto a 6-well plate one day before transfection. On the next day, the cells were transfected with 500 ng pB-TRE3G-NLS_SV40_-dCas9-3X NLS_SV40_-24X GCN4_-V4_-NLS_SV40_-P2A-BFP-PuroR-P2A-rtTA, 500 ng pB-TRE3G-ScFV-sfGFP-GB1-NLS_SV40_-HygroR-P2A-rtTA, and 200 ng pCAG-hyPBase using Chemifect. After 48 hr, cells were subjected to hygromycin (200 μg/ml) and puromycin (5 μg/ml) selection. After incubation for two weeks, cells with appropriate expression level of BFP and GFP were selected using FACS. Single cell clones were harvested for imaging a month later.

### Immunofluorescence

Cells were grown on 35mm glass bottom dish. After the coverage of cells reached 60-70%, cells were fixed with 4% PFA for 15 min, permeabilized with 0.5% Triton in PBS for 5 min and then blocked in blocking buffer containing 5% BSA and 0.1% Triton for 30 min. The cells were then incubated with primary antibodies in blocking buffer for 1 hr at room temperature, washed with PBS three times, and then stained with organic dyes-labeled secondary antibodies in blocking buffer for 1 hr at room temperature. The labeled cells were washed again with PBS, then post-fixed with 4% PFA for 10 min and finally stained with DAPI (Invitrogen).

Primary antibodies used in this study were lamin A/C (ab40467, Abcam, dilution 1:200), lamin B1 (sc6216, Santa Cruz, dilution 1:200). Secondary antibodies were donkey anti-rabbit Alexa Fluor 555 (A-31572, Thermo Fisher Scientific), donkey anti-goat Cy5 (705-005-147, Jackson Immuno Research Laboratories, dilution 1:50), donkey anti-mouse Cy3b (715-005-151, Jackson Immuno Research Laboratories, dilution 1:50).

### Optical setup and image acquisition

Briefly, all tracking experiments in living cells were performed on an Olympus IX83 inverted microscope equipped with a 100× UPlanSApo, N.A. 1.40, oil-immersion phase objective and EMCCD (DU-897U-CS0-#BV). The microscope stage incubation chamber was maintained at 37 °C and 5% CO_2_. A 488-nm laser (2RU-VFL-P-300-488-B1R; MPB) was used to excite the sfGFP fluorophore. The laser power was modulated by anacousto-optic-tunable-filtre (AOTF) and the beam width was expanded fivefold and focused at the back focal plane of the objective. The power density at the sample, with epifluorescence illumination, was 10 µW at 488nm. The microscope was controlled by home-written scripts. Movies of chromatin dynamics in living cells were acquired at 10 Hz. The motions of loci were studied by recording their trajectories in 2D rather than in 3D to increase time resolution and reduce phototoxicity. According to previous study, there is no significant difference in the movement volumes and diffusion coefficient of telomeres between different cell cycle stages in interphase, thus we collected images in interphase without further distinguishing between sub-stages of interphase.

For fixed cell conventional imaging experiments, an UltraVIEW VoX spinning disc microscope (PerkinElmer) was used. STORM imaging of lamin B1 was done on N-STORM (Nikon, Japan)

### Image analysis

All image stacks were analyzed using MATLAB tracking package ‘U-track’. Fluorescent puncta were identified in each frame with 2D Gaussian fitting after Fourier low-pass filtering. The coordinates of the fluorescent puncta were determined. Trajectories were created by linking identified puncta to their nearest neighbor within a maximum distance range of 5 camera pixels (800 nm) in the previous frame. Particles with trajectory gap larger than 10 consecutive frames were treated as two particles.

For each trajectory, the mean square displacement (MSD) as a function of time delay was calculated by the following equation:

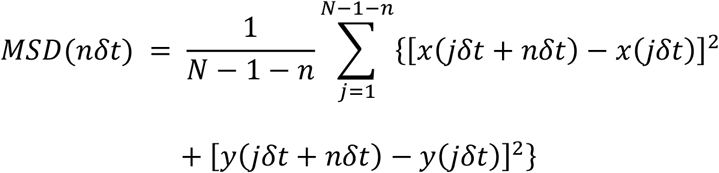

where δt is the time interval between two successive frames, x(t) and y(t) are the coordinates at time t, N is the total number of frames, and n is the number of time intervals. To maximize precision in long-range MSD, intervals smaller than N/10 were used for the calculation.

The analysis of MSD curves was carried out using custom MATLAB scripts. Each individual MSD curve was fitted by least-squares regression to the following model:

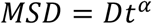

where D is the diffusion coefficient and α is the scaling factor. For each repeat, many trajectories were fitted and grouped. Additionally, every collected cell was inspected carefully, and any cell with slight motion was discarded to eliminate the contamination of such drift in the analysis of loci trajectories.

The three-dimensional image analysis was carried out in Imaris (Bitplane) by ImarisCell, a module designed specifically to identify, segment, track, measure and analyze cell, nucleus and vesicles in 3D images. Using “Cell Boundary from Cytoplasm” function, the nucleus was segmented by DAPI channel as the nuclear boundary and the genomic loci were fitted with 3D ellipsoid function as a spot. Then the shortest distance between the spot and the surface was calculated. For chromosome painting image analysis, “Surface” function was used to segment nuclear boundary by DAPI channel and territory boundary of chromosome 2 and chromosome 18 by 488 nm and 561 nm channel intensity. The volume and center of mass of nucleus and chromosome territories were output directly. The volume of each nucleus was measured to normalize the volume of chromosome territories. Distance between nuclear center of mass and chromosome territories was normalized by the cubic root of nuclear volume.

The threshold for subnuclear position assignment of loci was as follows: 4 pixels’ distance between the locus and nuclear envelope (640 nm), referring to previous publication about LMNB1 LAD FISH analysis (defined there as < 700nm, or 8 pixels, from the nucleus edge)(63).

STORM original data was processed by Insight3, ImageJ, and finally reconstructed to an image by home-written MATLAB scripts(64).

### Western blot

The cell lysates were blotted against the following primary antibodies: lamin B1 (sc6216, Santa Cruz, dilution 1:500) and β-actin (sc47778, Santa Cruz, dilution 1:500). The blots were visualized with peroxidase-coupled secondary antibodies.

### PI staining

Cells grown on 60 mm dish were digested by trypsin and collected to 1.5 ml tube. After being washed with PBS twice, cells were fixed in pre-chilled 75% ethanol at −20 °C overnight. The fixed cells were washed to remove ethanol, and then incubated in solution of 100 µg/ml RNase A and 0.2 % Triton X-100 for 30 min at 37 °C. Subsequent centrifugation of the samples was followed by a wash in PBS and staining with PI solution (50 µg/ml PI, 0.2% Triton X-100) at room temperature for 30 min. Cells stained with PI were analyzed in Flow cytometer (BD LSRFortessa^TM^) directly(65).

### Chromosome painting

Cells were grown on 22 × 22 mm^2^ coverslips. After the coverage of cells reached 70-80%, cells were fixed at −20 °C for 20 min in a pre-chilled solution of methanol and acetic acid at 3:1 ratio and then treated with 10 µl of probe mix with 5µl of each probe. The probe mix immersed cells were covered with a glass slide (25 × 75 mm) and sealed with rubber cement. The sample and probe were denatured simultaneously by heating slide on a hotplate at 75 °C for 2 min and incubated in a humidified chamber at 37 °C overnight. The coverslip was removed carefully from slide, washed in 0.4 × SSC at 72 °C for 2 min, and then in 2 × SSC, 0.05% Tween-20 at room temperature for 30 seconds. The labeled cells were rinsed briefly in PBS and finally mounted with ProLong® Diamond Antifade Mountant with DAPI (P36962, Thermo Fisher Scientific).

Chromosome 2 painting probe mix was XCP 2 green (D-0302-100-FI XCP 2, Metasystems) and Chromosome 18 painting probe mix was XCP 18 orange (D-0318-100-OR XCP 18, Metasystems).

### Hi-C experiment

Hi-C experiment was performed following the in situ Hi-C protocol(46). Briefly, cells were grown to about 70-80 % confluence, washed with PBS, crosslinked with 1%, v/v formaldehyde solution, and the reaction was quenched by 0.2M glycine solution. Cells were lysed and DNA was then cut with MboI and the overhangs were filled with a biotinylated base. Free ends were then ligated together in situ. Crosslinks were reversed, the DNA was sheared to 300-500bp and then biotinylated ligation junctions were recovered with streptavidin beads.

Sequencing libraries were generated using standard Illumina library construction protocol. Briefly, ends of sheared DNA were repaired and the blunt ends were added an “A” base to ligate with Illumina’s adapters that have a single ‘T’ base overhang. Then DNA was PCR amplified for 8-12 cycles. At last, products were purified using AMPure XP system and sequenced through XTen (Illumina).

### Hi-C data analysis

Hi-C data analysis was performed with HiC-Pro(66). Briefly, reads were first aligned on the hg19 reference genome. Uniquely mapped reads were assigned to restriction fragments. Then the invalid ligation products were filtered out, and eligible read pairs were counted to build Hi-C contact maps. At last, ICE normalization(67) was used to normalize the raw counts data.

#### Compartment A/B analysis

ICE-normalized 500-kb resolution matrices were used to detect chromatin compartments by R package HiTC(68). The whole genome was divided into two compartments based on the positive or negative values of the first principal component. The part with higher gene density was assigned as compartment A and the other part as compartment B.

#### TAD analysis

ICE-normalized 40-kb resolution matrices were used to detect TAD by Perl script matrix2insulation.pl (http://github.com/blajoie/crane-nature-2015). Insulation scores were calculated for each 40-kb bin, and the valleys of insulation score curves were defined as TAD boundaries. TADs smaller than 200 kb were filtered out as in previous method(51).

## General

We thank Dr. Ronald D. Vale (University of California, San Francisco) for providing SunTag plasmids, Dr. Wensheng Wei (School of life sciences, Peking University) for providing sgRNA expression plasmid, Dr. Feng Zhang (Broad Institute) for providing plasmids px330 (Addgene Plasmid # 42230) and Dr. Wei Guo (Department of Biology, University of Pennsylvania) for providing the MDA-MB-231 cell line. We also thank Dr Hongxia Lv at the core imaging facility of the School of Life Sciences, Peking University for imaging support.

## Author contributions

Y.S., C.L., L.C., and S.S. conceived and designed the experiments. L.C. performed all the cloning, immunofluorescence, western blot, chromosome painting, live cell tracking experiments and image data analysis. M.L. performed Hi-C experiments and conducted data analysis of Hi-C and RNA-seq. S.S. wrote the MATLAB script for images analysis and carried out the stable cell line construction and knockout experiments. Y.H. gave help in FISH experiments. B.X. involved in the critical discussion. L.C., M.L., S.S., C.L., and Y.S. wrote the manuscript.

## Supplementary Figures

**Fig. S1.**
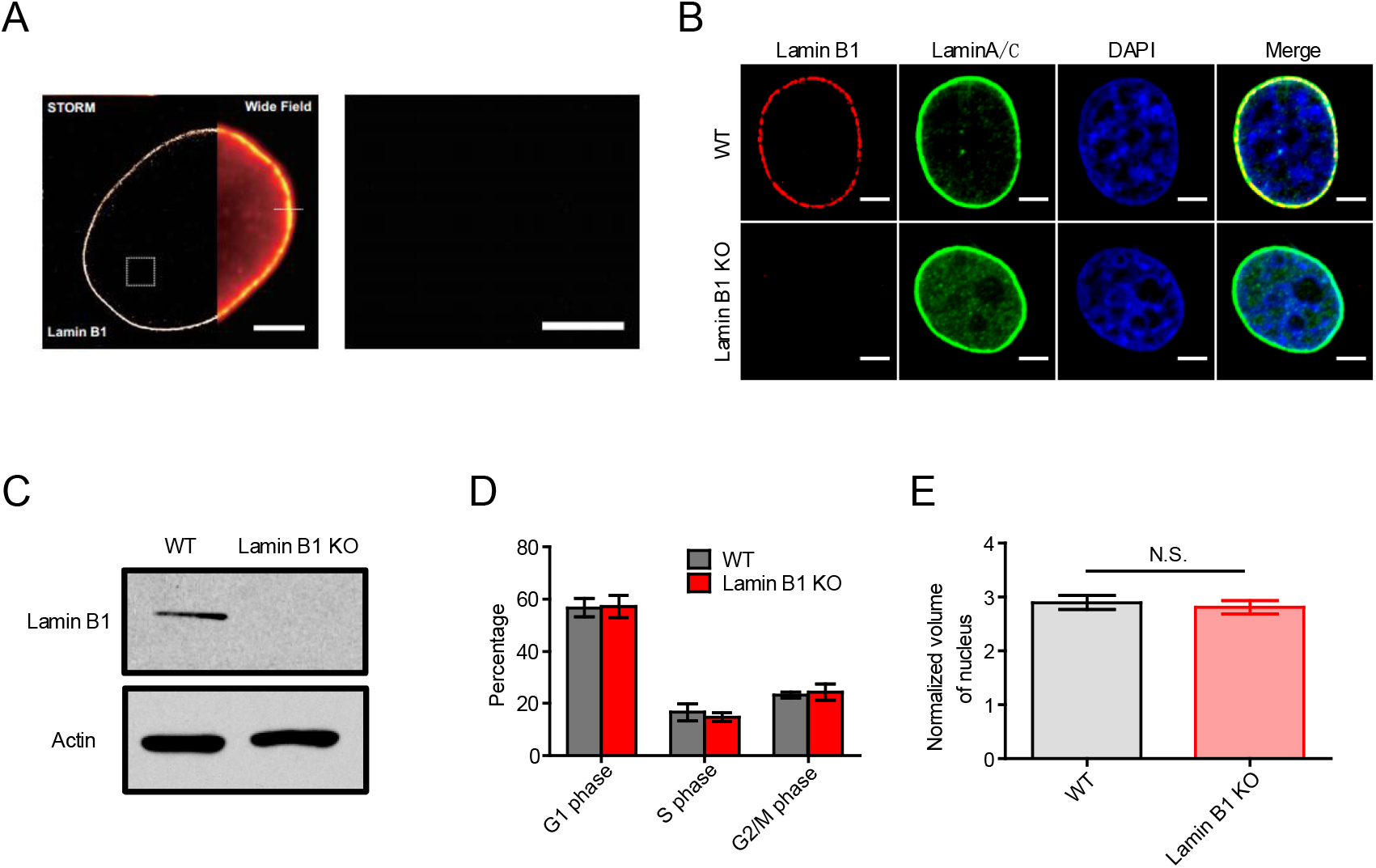
Preparation of lamin B1 knock-out (KO) cell lines. (A) STORM image and wide-field image of lamin B1 in MDA-MB-231 cells. Scale bar: 5 μm. Magnified image of the boxed area shows that lamin B1 almost has no presence in nucleoplasm. Scale bar: 1 μm.
(B) Immunofluorescence of chosen lamin B1-KO clone with lamin B1 and lamin A/C antibody. WT MDA-MB-231 cell line is used as positive control. The result indicates that lamin B1 is totally knocked out in the chosen KO clone, while lamin A/C is unaffected. The images are shown under the same intensity threshold between WT and lamin B1-KO cells.
(C) Western blot of chosen lamin B1 KO clone with lamin B1 antibody. Actin is used as loading control. WT MDA-MB-231 cell line functions as positive control.
(D) Percentage of cells in G1, S, and G2/M phase in 3 independent experiments.
(E) Quantification of nuclear volume based on DAPI staining in WT (n=18) and lamin B1-KO (n=16) cells. Mean ± SE. Nuclear volumes of the two samples do not have significant difference.

**Fig. S2.**
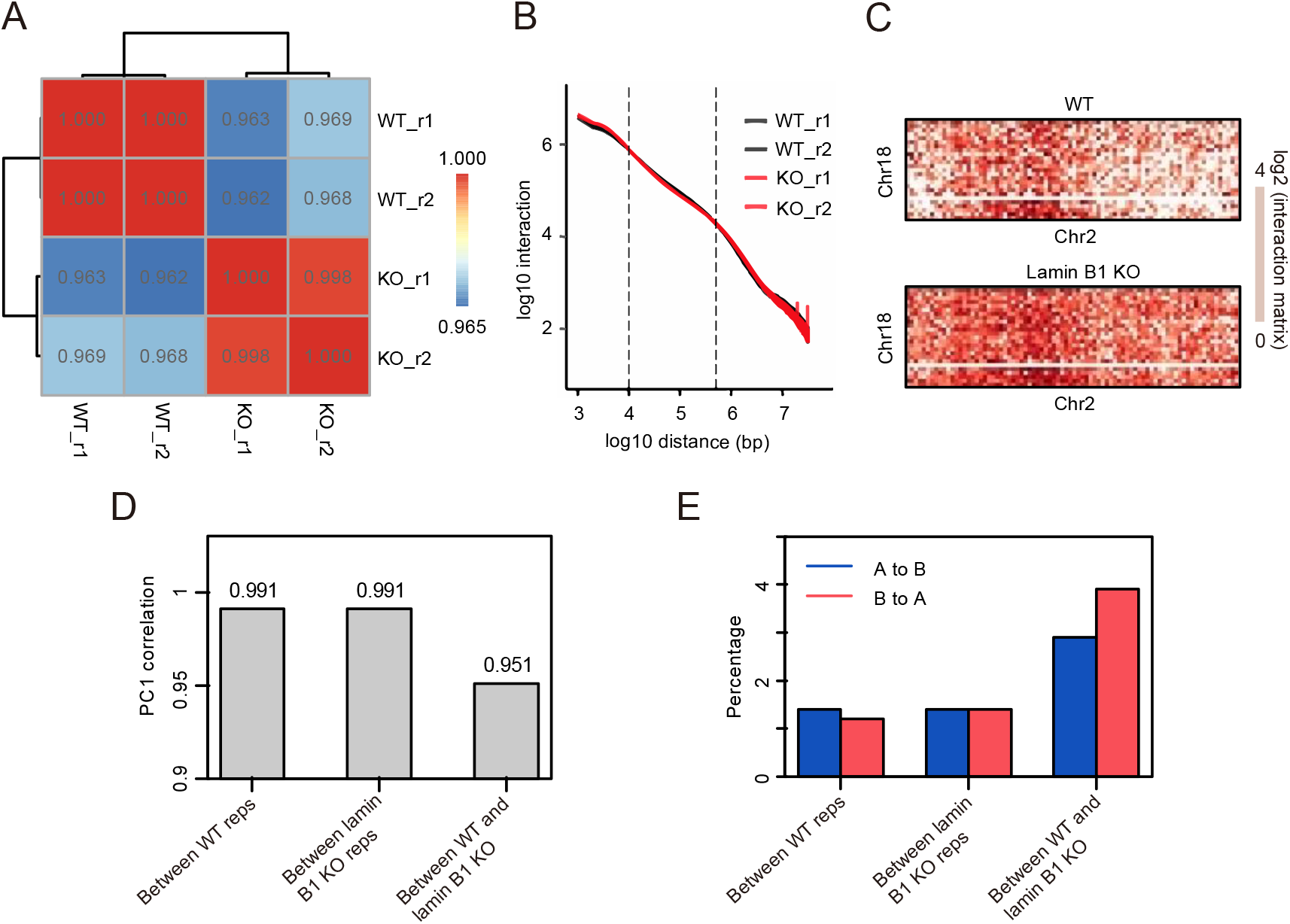
Reproducibility analysis of Hi-C data. (A) Pearson correlation coefficients of the whole genome interaction matrices (resolution: 500 kb) of WT and lamin B1-KO replicates.
(B) Hi-C interaction frequency as a function of genomic linear distance for WT and lamin B1-KO replicates.
(C) Normalized Hi-C trans-interaction matrices for chromosome 2 and 18 in WT and lamin B1-KO samples.
(D) Pearson correlation coefficients of the whole genome PC1 values between WT replicates (r=0.991), lamin B1-KO replicates (r=0.991), WT and lamin B1-KO samples (r=0.951).
(E) Percentage of the whole genome A/B compartment transition between WT replicates, lamin B1-KO replicates, WT and lamin B1-KO samples.

**Fig. S3.**
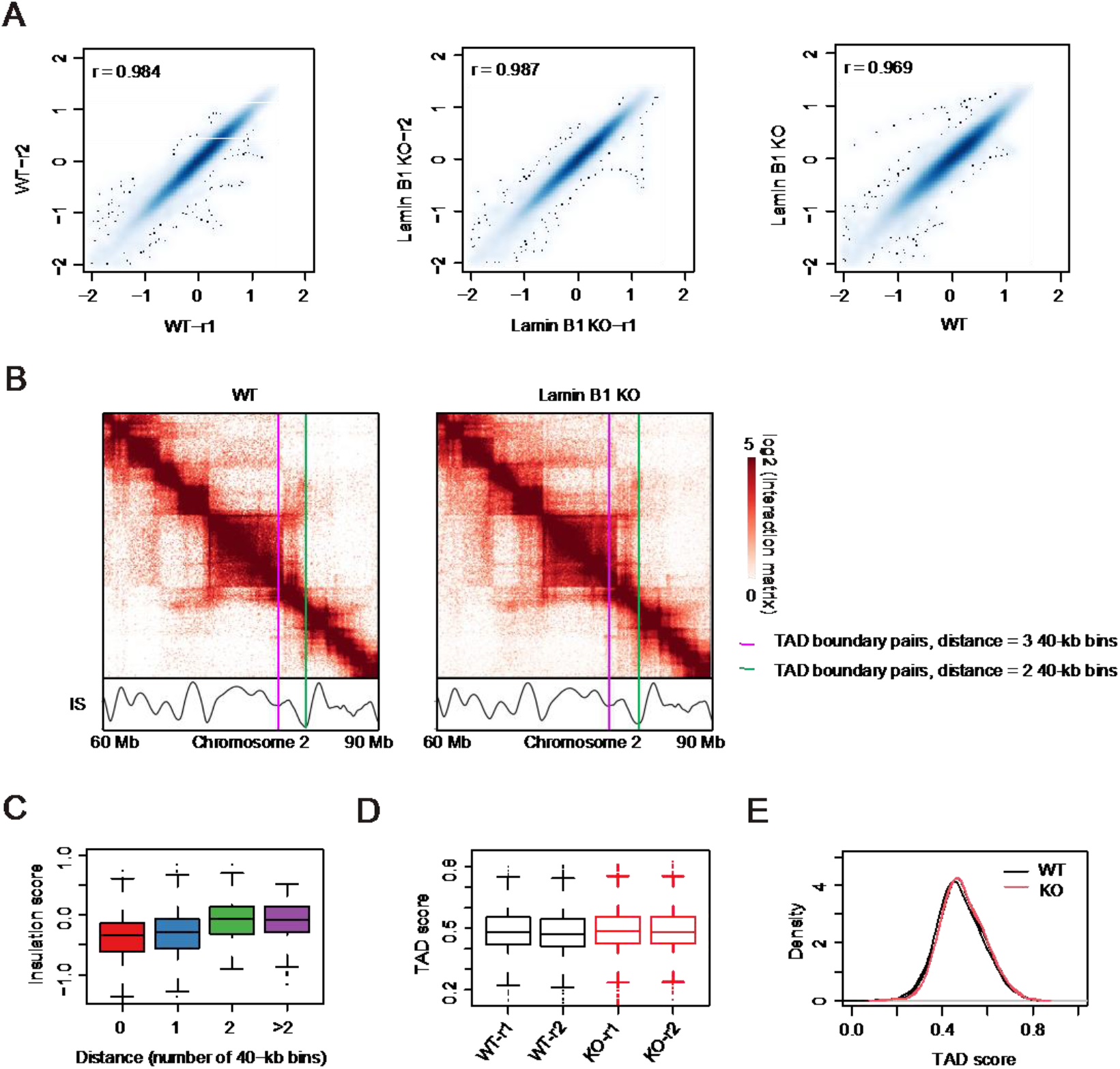
TAD analysis. (A) Scatter plots of the whole genome insulation scores of WT replicates (left, Pearson correlation coefficient r=0.984), lamin B1-KO replicates (middle, Pearson correlation coefficient r=0.987), WT and lamin B1-KO samples (right, Pearson correlation coefficient r=0.969).
(B) Example of two most adjacent TAD boundary pairs in WT and lamin B1 KO-cells, with the distance of 3 (purple) and 2 (green) 40-kb bins.
(C) Boxplot of mean insulation scores of most adjacent TAD boundary pairs in WT and lamin B1-KO cells at a distance of indicated numbers. Together with Fig. S3B, the insulation score valleys at identical boundary pairs are lower and sharper than the changed ones, indicating that shifted TAD boundaries are due to variance upon calculation.
(D) Boxplot of TAD scores of WT and lamin B1-KO replicates. TAD score=intra-TAD interactions / (intra- + inter-TAD interactions).
(E) TAD score distribution of WT and lamin B1-KO samples.

**Fig. S4.**
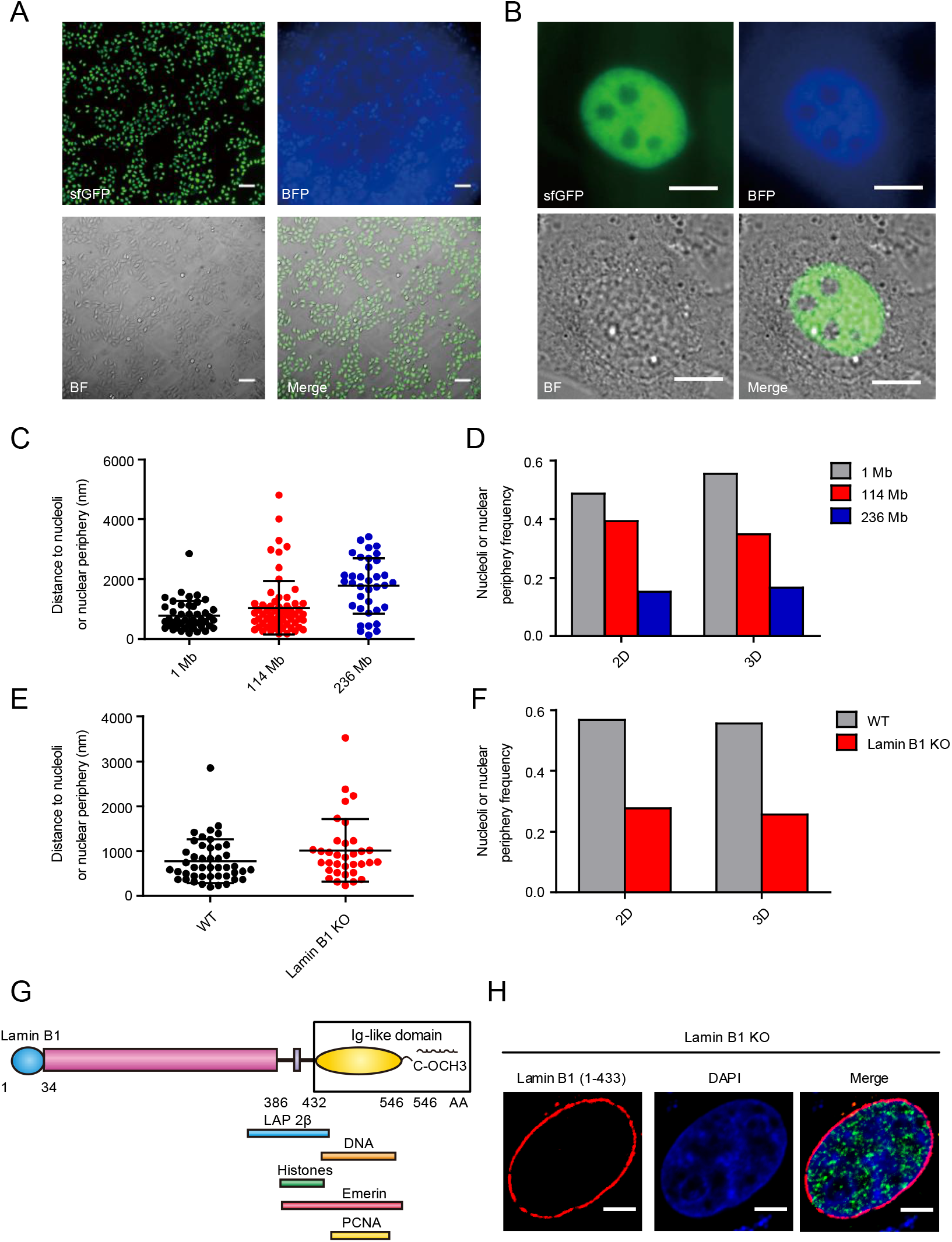
Construction of SunTag stable cell line, comparison between 2D and 3D images and description of lamin B1 truncation. (A) Fluorescent images of the SunTag cell line under 10X magnification with Dox induction. BFP and bright field (BF) are also shown. All cells display similar expression level of both dCas9-(GCN4) X24 and scFv-GCN4-sfGFP. Scale bars: 100 μm.
(B) Fluorescent images of the SunTag cell line under 100X magnification with Dox induction. BFP and bright field (BF) are also shown. The absence of fluorescent signal of both blue and green channel in nucleoli indicates that this method is superior to the conventional dCas9-GFP labeling method which shows severe nucleoli aggregation. Scale bars: 10 μm.
(C) Quantification of distance from three genomic loci on chromosome 2 to nucleolus or nuclear periphery in reconstructed 3D images. 2 independent experiments.
(D) Nucleoli and nuclear periphery localization frequency of 3 genomic loci on chromosome 2 from 2D and 3D images. Each locus is assigned to this localization according to the rule that the minimum distance to nucleoli or nuclear envelope is less than 4 pixels (∼ 640 nm).
(E) Quantification of distance from 1 Mb locus on chromosome 2 to nucleolus or nuclear periphery in WT and lamin B1-KO cells in reconstructed 3D images. 2 biological repeats.
(F) Nucleoli and nuclear periphery localization frequency of 1 Mb locus on chromosome 2 from 2D and 3D images. Each locus is assigned to this localization according to the rule that the minimum distance to nucleoli or nuclear envelope is less than 4 pixels (∼ 640 nm).
(G) Diagram of lamin B1 and binding sites of interaction partners. The Ig-like domain of lamin B1 mediates direct and indirect interaction between lamin B1 and chromatin through DNA and other proteins.
(H) Lamin B1 (1-433) truncation without Ig-like domain localizes in nuclear periphery when expressed in lamin B1-KO cells.

**Fig. S5.**
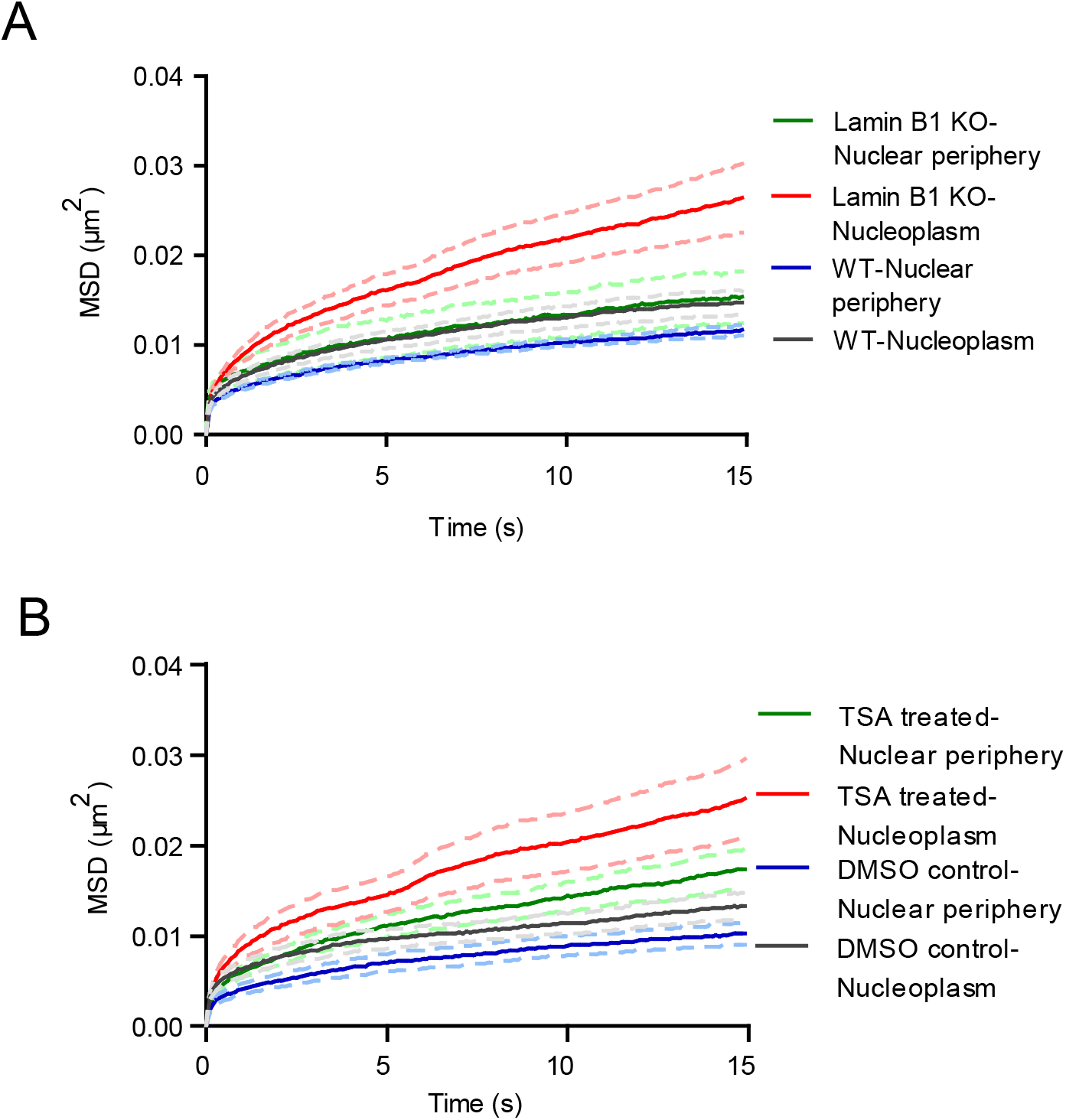
MSD curves of 1Mb loci on chromosome 2. (A) SD curves of 1Mb loci on chromosome 2 localized in nuclear periphery and nucleoplasm of WT and lamin B1-KO cells, separately. Mean ± SEM. Depletion of lamin B1 increases 1Mb loci dynamics in both periphery and nucleoplasm. 3 independent experiments.
(B) MSD curves of 1Mb loci on chromosome 2 localized in nuclear periphery and nucleoplasm of DMSO-treated control and TSA-treated cells, separately. Mean ± SEM. TSA treatment increases 1Mb loci dynamics in both nuclear periphery and nucleoplasm. 3 independent experiments.

**Fig. S6.**
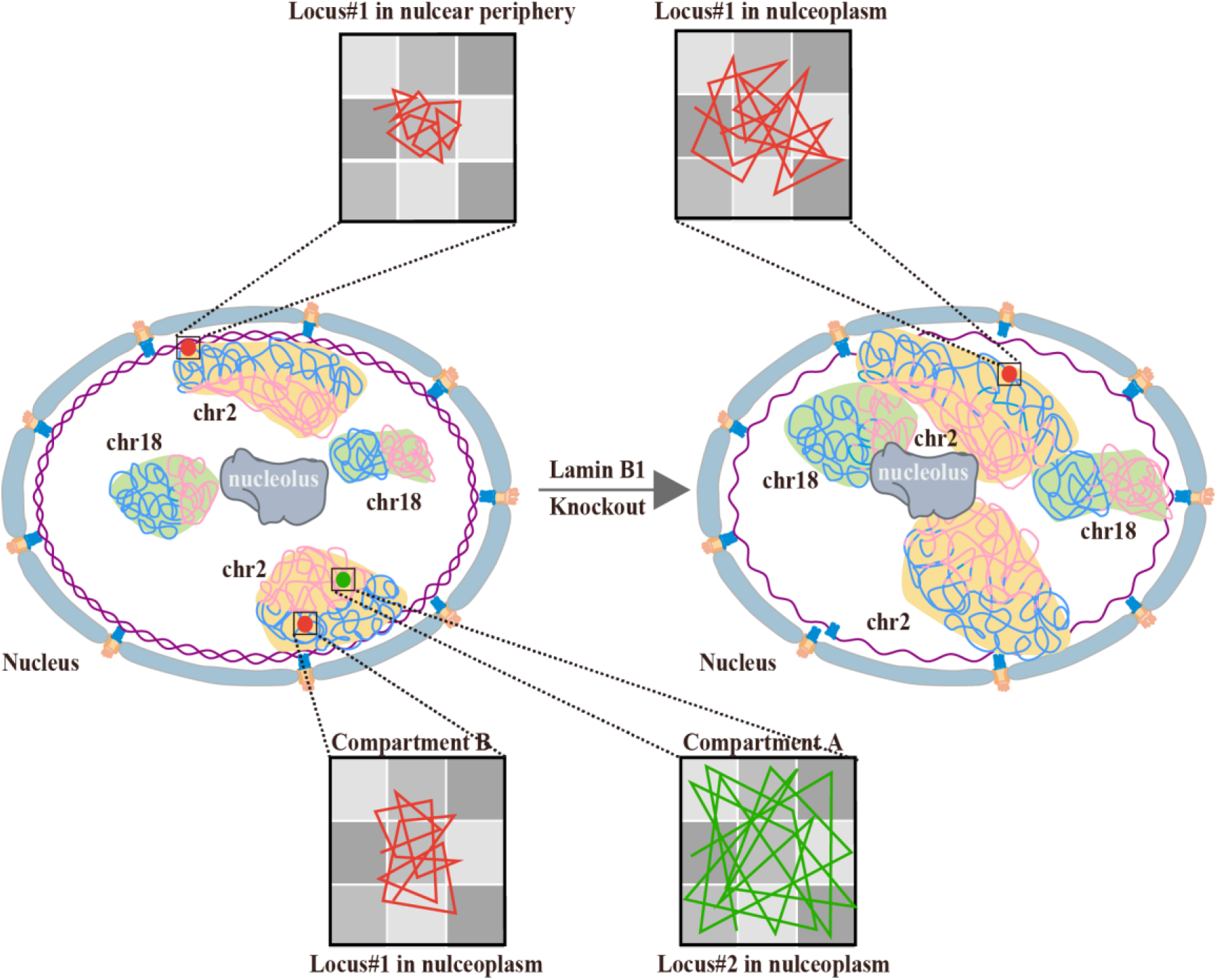
Model. A model describing lamin B1 regulating chromatin higher-order structure and dynamics through tethering to chromatin. Loss of lamin B1 in lamina releases a part of chromatin from nuclear periphery to nuclear interior. The change of chromosome compaction state induces expansion of chromosome territories and thus increases the interaction ratio between different chromosomes. Besides, loss of lamin B1 reduces the integrity and segregation of chromatin compartments and part of genomic regions switches between A and B compartments. However, lamin B1 is not required for TAD insulation. Furthermore, depletion of lamin B1 can increase genomic loci dynamics. The dynamic motion of the same locus in different subnuclear regions demonstrates significant difference, and nuclear periphery-localized loci is much less mobile than the nucleoplasmic-positioned loci. Besides, chromatin compaction is a more fundamental factor affecting chromatin dynamics. Genomic loci in less compact compartment A are of higher mobility than those in more compact compartment B.

**Fig. S7.**
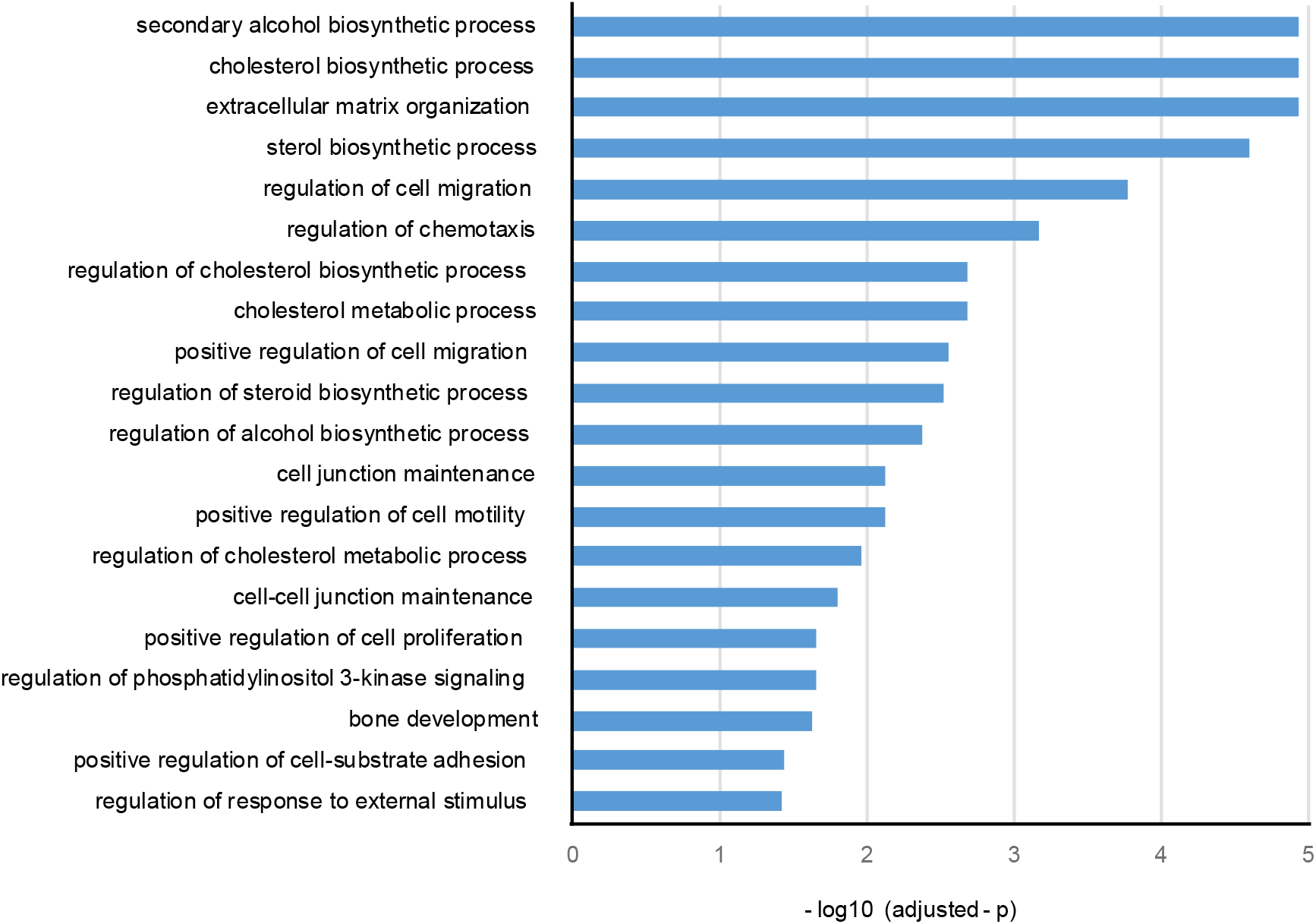
GO (Biological Process) analysis of down-regulated genes upon lamin B1 KO.

**Table S1.** Quality control statistics for Hi-C data processing.

|  | WT-r1 | WT-r2 | Lamin B1 KO-r1 | Lamin B1 KO-r2 |
| --- | --- | --- | --- | --- |
| **Total read pairs<sup>[1]</sup>** | 138,193,618 | 223,885,207 | 250,362,318 | 253,024,668 |
| **Uniquely aligned read pairs<sup>[2]</sup>** | 108,155,179 | 172,561,205 | 182,642,642 | 194,536,722 |
|  | 78.26% | 77.08% | 72.95% | 76.88% |
| **Valid interaction<sup>[3]</sup>** | 95,264,155 | 152,469,233 | 148,123,567 | 166,884,723 |
|  | 88.08% | 88.36% | 81.10% | 85.79% |
| **Self-Circle<sup>[4]</sup>** | 123,564 | 178,371 | 520,046 | 222,818 |
|  | 0.11% | 0.10% | 0.28% | 0.11% |
| **Dangling-end<sup>[5]</sup>** | 2,726,851 | 4,213,273 | 13,818,685 | 6,521,973 |
|  | 2.52% | 2.44% | 7.57% | 3.35% |
| **Valid interaction rmdup<sup>[6]</sup>** | 79,041,927 | 131,026,015 | 129,693,005 | 134,947,244 |
|  | 73.08% | 75.93% | 71.01% | 69.37% |
| **Trans_interaction<sup>[7]</sup>** | 7,834,781 | 13,603,317 | 15,747,086 | 16,307,003 |
|  | 9.91% | 10.38% | 12.14% | 12.08% |
| **Cis_interaction<sup>[8]</sup>** | 71,207,146 | 117,422,698 | 113,945,919 | 118,640,241 |
|  | 90.09% | 89.62% | 87.86% | 87.92% |
| **Cis_shortRange<sup>[9]</sup>** | 22,113,855 | 34,770,742 | 37,869,246 | 39,050,597 |
|  | 31.06% | 29.61% | 33.23% | 32.92% |
| **Cis_longRange<sup>[10]</sup>** | 49,093,291 | 82,651,956 | 76,076,673 | 79,589,644 |
|  | 68.94% | 70.39% | 66.77% | 67.08% |

The percentage denominators of [2] are the read pair numbers in [1]; the percentage denominators of [3][4][5][6] are the uniquely aligned read pair numbers in [2]; the percentage denominators of [7][8] are in [6]; and the percentage denominators of [9][10] are in [8].

Movie. A representative movie and tracking trajectory of labeled genomic locus in a living cell.

